# A Deep Learning Approach for Modelling the Complex Relationship between Environmental Factors and Biological Features

**DOI:** 10.1101/2023.06.26.546510

**Authors:** Devashish Tripathi, Analabha Basu

**Affiliations:** National Institute of Biomedical Genomics, Kalyani 741251 West Bengal, India

**Keywords:** deep learning, genomics, natural selection

## Abstract

Environmental factors play a pivotal role in shaping the genetic and phenotypic diversity among organisms. Understanding the influence of the environment on a biological phenomenon is essential for deciphering the mechanisms resulting in trait differences among organisms. In this study, we present a novel approach utilizing an Artificial Neural Network (ANN) model to investigate the impact of environmental factors on a wide range of biological phenomena. Our proposed workflow includes hyperparameter optimization using model-based methods such as Bayesian and direct-search methods such as Random Search, and a new approach combining random search and linear models (RandomSearch+lm) to ensure a robust ANN architecture. Moreover, we employed a generalized version of the variable importance method to generate the feature importance metric using estimated weights from ANN. By applying this comprehensive ANN-based approach to functional genomics, we can gain valuable insights into the mechanisms underlying trait differentiation in organisms, while simultaneously enabling prediction and feature selection tasks. This methodology provides a robust and efficient framework for studying the complex relationships between environmental factors and biological features in biological systems.

## Introduction

The role of environmental factors in shaping the genetic and phenotypic diversity among organisms has been a central question in genomic research. It is well acknowledged that the complex interaction between the environment and organisms plays a crucial role not only in the evolution of biological functions in response to environmental factors but also in the modification of biological functions throughout an organism’s life cycle. Understanding how environmental factors influence the genetic composition and phenotypic traits of organisms is essential to unravel the mechanisms underlying adaptations through natural selection and phenotypic diversity among organisms.(1,2,3)

Researchers have investigated a wide range of biological phenomena influenced by environmental factors. Adaptation of populations to their local environment, known as local adaptation, is one such phenomenon. In quantitative genetics, a phenotype can be expressed as a function of genotype, environment, and genotype-environment interactions (G × E). The G × E component contributes to the evolution of quantitative traits, and thus trait differences within and among populations structured in a local geographical space. Since polygenic traits are a function of multiple quantitative trait loci (QTL), the adaptation of populations along an environmental gradient results in allele frequency patterns associated with environmental factors. In functional genomics, it has been well established that the environment influences gene expression, gene regulation, and eventually the phenotype throughout an organism’s life cycle. Gene expression is a biological process of decoding information from DNA through transcription and translation. The process eventually produces a protein or noncoding RNA, which ultimately affects the phenotype of the organism. Gene regulation is a complex mechanism through which cells control the time and amount of expression of specific genes. Thus, investigating the influence of environmental factors on gene expression and gene regulation can provide valuable insights into the mechanisms underlying trait differentiation in organisms(12,13,14).

Researchers have implemented a diverse set of mathematical models, ranging from statistical linear models to deep learning methods, to decipher genetic variants that show strong evidence of association with environmental factors. One major purpose of these methods is to yield a probable set of genetic variants under natural selection. Statistical linear models assume a linear relationship between the genotype and environmental factors, which may not necessarily be true in a real geographical landscape. Allele frequencies may not change linearly across geographical space and evolutionary forces like natural selection will give rise to complex patterns which linear models are unable to capture. (4,5,6,7)

Deep learning models have been implemented in this scenario, as they can learn complex relationships between genotypes and the environment (8,9). To implement deep learning models, researchers have used the genotype matrix as the input data and environmental factors as the output data. In deriving the architecture of neural networks, random search and grid search, with a limited number of trials are implemented for basic hyperparameter optimization. We demonstrate that the derived architecture might not be robust because it does not explore the large hyperparameter space during hyperparameter optimization. The implemented workflow is also not an efficient approach, because if we assume ‘n’ environmental factors, it performs hyperparameter optimization for each pair of environmental factor and genotype matrix. This eventually scales the time linearly the increasing number of environmental factors (10). There are no restrictions, such as, defining the output layer with only one neuron in an Artificial Neural Network (ANN). An ANN can take ‘n’ neurons in the output layer and model the relationship between the input and output data. (11)

The attempt to model the relationship between biological features and the environment poses challenges due to its high dimensionality. We propose using the biological features as input data to identify the features influence by environment. This approach estimates the effects of individual biological features while accounting for the presence of other input biological features and can capture the interaction between the input features. Here, we propose an ANN for modelling a wide range of biological phenomena that are influenced by the environment. An ANN can detect biological features that are highly influenced by the environmental factors in a biological system. We are developing a model that can be generally used for prediction and feature selection problems. In summary, the proposed workflow requires an input matrix of biological features and an output matrix of environmental factors (output variables ≥ 1), performs hyperparameter optimization using Bayesian optimization, hyperband optimization, and a new RandomSearch+lm approach, trains the model with the optimized ANN architecture, and implements a generalized version of the variable importance method (27) to estimate the feature importance metric using estimated weights from all the layers in ANN.

## Methodology

### Artificial Neural Network

An ANN is a supervised deep learning model that learns the complex relationship between input data and output variables. In the proposed framework, the input data are the biological features matrix, and the output data are environmental factors. The general architecture of a neural network consists of an input layer, an output layer, and a set of hidden layers, as shown in the figure below. The input layer takes an input vector of the *p* biological features and the output layer has *k* environmental factors corresponding to each input vector.

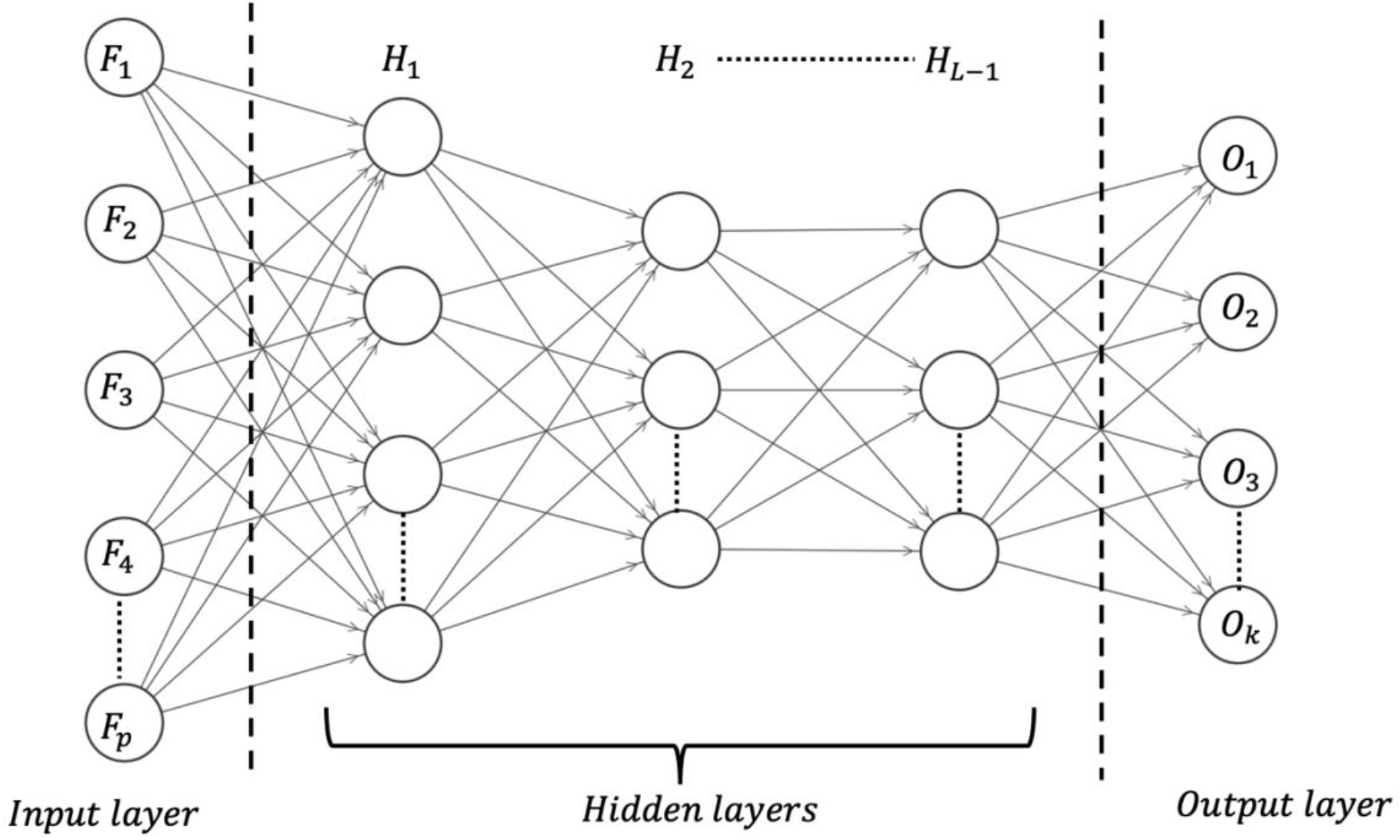

### General Architecture

We define the components of the ANN as shown in the figure above. In this figure, we do not explicitly show the bias units corresponding to each layer. The contribution of the bias unit can be seen in the mathematical derivations.

*L* = *Total number of layers in Neural Network*,

*a*^*l*^ *= output vector from layer l*,

*a*^*L*^ *= predicted output vector from final layer*,

*n*^*l*^ *= number of neurons in l*^*th*^ *layer*,

*ϕ*^*l*^ *= activation function for l*^*th*^ *layer*,

*b*^*l*^ *= bias vector for l*^*th*^ *layer*,

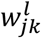 *=weight for j*^*th*^ *neuron in layer l receiving input from k*^*th*^ *neuron in layer l* − 1

The weights between two layers (*l, l* − 1) can be represented in a matrix form as follows

*k neurons in layer l* − 1

*W*^*l*^ = *Weight matrix for layer l* = 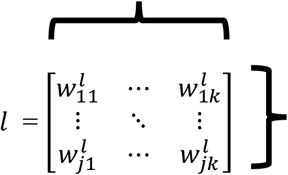 *j neurons in layer l*

In general, the output from any layer *l* can defined as below

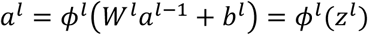

The predicted output from the final layer becomes a composite function of the outputs from all previous layers.

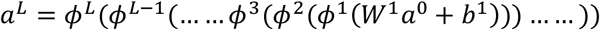

### Training ANN

To train the ANN, feedforward and backpropagation approaches are used to estimate the weights and biases of the ANN such that the cost function reaches an optimum.

### Feedforward

In the feedforward approach, weights and biases are initialized across the layers in the ANN, training examples are then processed through the network, and the final output is predicted. A schematic process is shown below

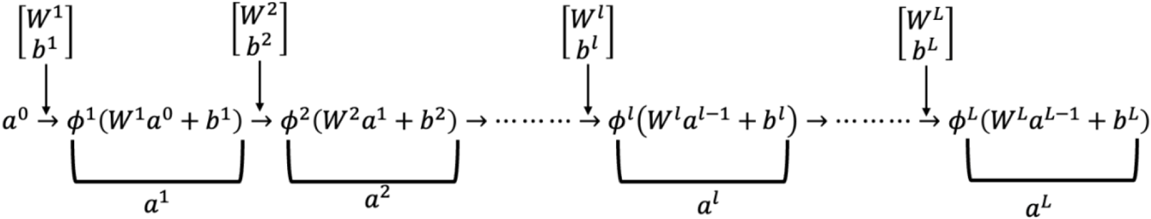

### Backpropagation

The predicted output from the final layer after the feedforward is compared with the actual output and the cost function is estimated. The error is backpropagated to minimize the cost function by updating the weights across the layers in ANN. A schematic process is shown below,

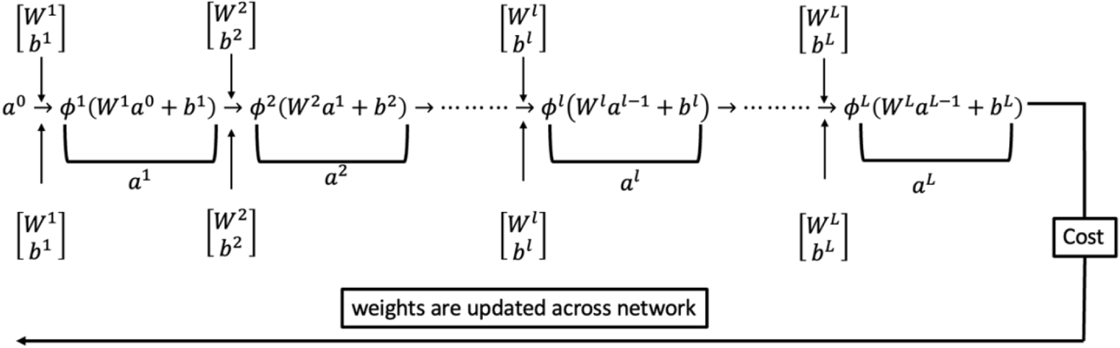

To perform the backpropagation the following four equations have been defined in the literature

1. error of all neurons in the last layer

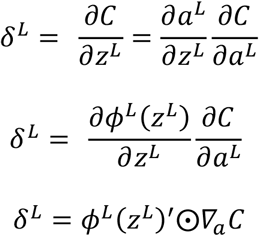
2. error of all neurons in layer *l*

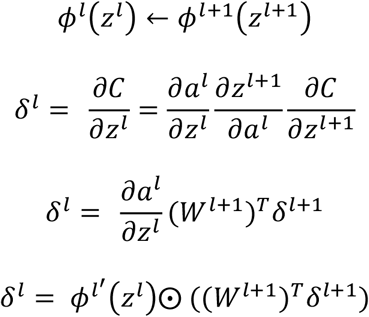
3. Derivative of cost with respect to bias in layer *l*

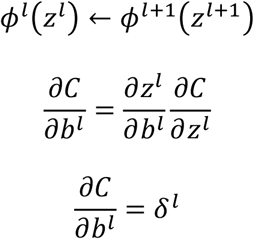
4. Derivative of cost with respect to weights in layer *l*

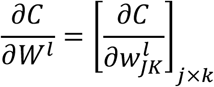

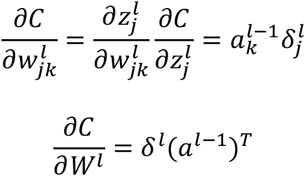

Where C is the cost function and can be chosen based on the research problem and the nature of the output variables. (16,17,18,19)

Assuming output variables are continuous in nature, the mean squared error cost function for a single training example can be defined as

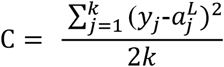; *y*_j_ *= actual output*, 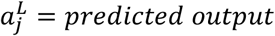 = *predicted output f or j^th^ neuron*

*n = size of training dataset*

*k = size of output layer*

The training approach discussed above can be implemented using the gradient descent optimization method (20). Here, we assume that ANN architecture has been fixed.

*Require*: *Initialization of weights and biases for each layer in ANN*

*Require*: *m*: *size of training examples batch*

*Require: n_epoch* : *number of epochs for training the ANN*

*FOR epoch =* 1 *to n_epoch*

*FOR i= 1 to m*

*Set the corresponding activation a*^*0*^ *for the input layer*// Input

FOR *l* = 1 *to L*

*compute and store z*^*l*^←*W*^*l*^*a*^*l−*1^+*b*^*l*^, *a*^*l*^← *ϕ*^*l*^ (*z*^*l*^) // Feedforward

END FOR

*δ*^*L*^ *Compute the vector δ*^*L*^←*ϕ*^*L*^(*z*^*L*^)′⨀*∇*_a_ *C* // Output layer error

FOR *l =L* − 1 *to* 1

*compute* 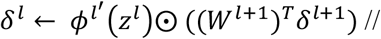 Backpropagate the error

*computeand store*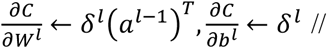Gradient of cost function with respect to *W*^*l*^, *b*^*l*^

END FOR

END FOR

*For l =L* − 1 *to* 1

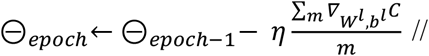 Update *W*^*l*^, *b*^*l*^ using mean of gradient of cost function

END FOR

END FOR

The above gradient descent algorithm (m = n) transforms into a stochastic gradient descent if m=1 and a mini-batch gradient descent if m<n.

The algorithm described above is slightly modified depending on the choice of optimization algorithms, such as Adam and RMSprop. These algorithms are extensions of gradient descent and implement the concept of adaptive learning rate optimization.

RMSProp (Root mean square propagation) modifies the parameters update equation by scaling the mean gradient by the square root of the exponentially weighted average of squared gradient and thus results in faster convergence to optima. (21)

Adaptive Moment Estimation (Adam) modifies the parameter update equation by replacing the mean gradient with the ratio of the exponentially weighted average of the gradient and the square root of the exponentially weighted average of the squared gradient. Adam provides an improvement over RMSProp as it incorporates information through a first-order moment of gradient as well. (22)

As we can see from the above protocol, we obtained estimates of parameter weights and biases from the data. In addition to these parameters, every other parameter, such as the number of hidden layers, number of neurons in each layer, activation function, optimization algorithm, and learning rate, are known as hyperparameters because they need to be set by the user and cannot be estimated using data. Thus, hyperparameter optimization has become a crucial part of deep learning architecture development.

### Hyperparameter optimization

In this section, we discuss current state-of-the-art methods and a new approach for hyperparameter optimization that merges random search and a statistical linear model.

Hyperparameter optimization can be described using a mathematical equation as follows

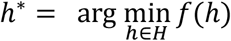

where, *f(h) = cost function to optimize such as MSE on validation set*

*h*^*^ *= hyperparameter configuration that optimizes f(h)*

### Random Search

The random search algorithm takes a random hyperparameter configuration (*h*_*i*_) from the parameter space. Perform the training of ANN by fixing the architecture based on *h*_*i*_, record the validation loss. Choose the *h*_*i*_ results in optimum loss after repeating the procedure for a fixed number of trials (23). Require: n_trial : number of trials

Require: H: hyperparameter space

SET i ←1

Draw a random sample of the hyperparameter set (*h*_1_) from *H*.

Fix neural network architecture based on *h*_1_

Train the fixed ANN and record the loss on the validation set SET Best_hyperparameters ← *h*_1_

FOR i = 2 to n_trial

Draw a random sample of hyperparameter set (*h*_*i*_) from *H*.

Fix neural network architecture on the basis of *h*_*i*_

Train the fixed ANN and record the loss on the validation set

IF *validation loss (h*_*i*_) < *validation loss (h*_*i*−1_) //minimising the loss function Best_hyperparameters ← *h*_*i*_

END IF

END FOR

There is always an uncertainty in choosing the number of trials required to reach the optimum validation loss as the number of trials required increases with the increasing size of the hyperparameter space.

### Hyperband

The hyperband further extends the random search algorithm. In summary, the idea behind the algorithm is that instead of training all the hyperparameter configurations (*h*_*i*_) for fixed number of epochs drop the worst performing *h*_*i*_ after reaching a threshold of epochs. In overview the algorithm’s objective is to find the best-performing *h*_*i*_ within a limiting budget such as the number of epochs, number of trials, and computational time (24).

### Bayesian optimization

Bayesian optimization is a model-based approach, which is a significant improvement over the direct-search algorithms described above. The idea behind Bayesian optimization is that instead of evaluating any random hyperparameter configuration (*h*_*i*_) first build a probabilistic model of the cost function (e.g., validation loss) given a *h*_*i*_. Sample *h*_*i*_ from the probabilistic model then perform the training of ANN, evaluate the cost function and update the probabilistic model. The algorithm implements a balanced exploration and exploitation approach i.e. sample *h*_*i*_ from most probable region as well as also explore less probable regions. The algorithm provides a robust architecture in a smaller number of trials than direct-search algorithms (25).

In general Bayesian optimization procedure requires the following components,

- The statistical distribution corresponding to each hyperparameter becomes the domain over which the best hyperparameter search must be performed.
- The cost function *f*(*h*) that has to be optimised on validation data
- A surrogate model of the cost function *f*(*h*)
- A selection function to take a decision of replacement of the current best hyperparameter configuration by *h*_*i*_
- Information from a pair of (validation loss, *h*_*i*_) is used by the algorithm to update the surrogate model

The choice of the surrogate model and selection function depends on the nature of the hyperparameter distributions and can be chosen based on the literature review. Some of the common choices for surrogate models are the Gaussian process, and Tree Parzen Estimators, and for selection function probability of improvement, expected improvement can be used.

### RandomSearch+lm

Here we present another model-based approach to search for the best hyperparameter configuration (*h*_*i*_) by merging both random search and linear model with first-order interaction. The Algorithm is described below

1. Using the random search algorithm generate a training dataset after N trials of the random search algorithm. In each trial *h*_*i*_and corresponding validation loss are recorded, the final data structure is shown below

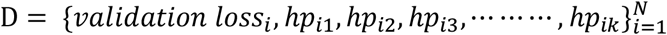
2. Following data generation, Now we have an input matrix of hyperparameters of size (*N* × *K*) and validation loss as output. We can fit a multiple linear regression model with first-order interactions to predict the validation loss given *h*_*i*_. This model can explain the impact of various hyperparameters on the validation loss as well as how the interaction among hyperparameters affects the validation loss. In general, a linear regression model with first-order interactions (26) can be defined as follows,

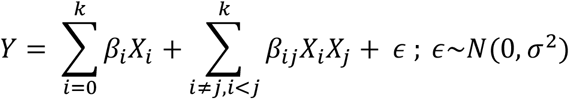

In this case, Y is validation loss and *X*_*i*_′*S* are hyperparameters.
3. After having a fitted model, we can process a large matrix of (*h*_*i*_)_*H*×*K*_ and predict the validation loss corresponding to each *h*_*i*_. The best *h*_*i*_ results in optimised predicted validation loss. Our approach can speed up the implementation of hyperparameter space search using a predictive model, without implementing a computationally expansive evolutionary algorithms which requires training the ANN from scratch for every choice of parameter set.

### Feature Selection

After hyperparameter optimization, we obtain the hyperparameter configuration that minimizes the validation loss and then the ANN architecture gets trained on the entire dataset.

After training the model, the next step is to estimate the contribution of each input feature to the output prediction. The contribution score for features helps select the most important features from the input set of features. The feature importance method (27) shows the mathematical formulation for measuring feature contributions in an ANN with one hidden layer and a single output in the output layer using the estimated weights from the ANN. The algorithm was implemented in the h2o package (28), which measures the contribution of the input feature in the final prediction using the estimated weights from the first two hidden layers. Here, we generalize this algorithm by incorporating weights from a finite number of hidden layers (*L – 1*) to measure contribution of input features in the final prediction of *K* outputs simultaneously in the output layer.

Let,

*n*_*h*_ *= number of neurons in h*^*th*^ *hidden layer*

*K = number of output neurons in output layer*

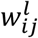*= weight for j*^*th*^ *neuron in layer l receiving input from i*^*th*^ *neuron in layer* (*l* – 1)

*N= number of input features*

*p*_*ij*_ = 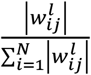

*p*_*ij*_ is defined as the contribution of *i*^*th*^ neuron in layer (*l* – 1) to the input of *j*^*th*^ neuron in layer *l*

Case 1: one hidden layer, one output layer

The contribution of *i*^*th*^ input feature in the prediction of *K*^*th*^ output can be written as 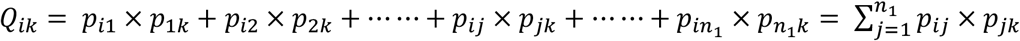

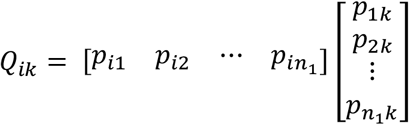

Thus the contribution of *i*^*th*^ input feature in the prediction of each of the *K* outputs can be written as

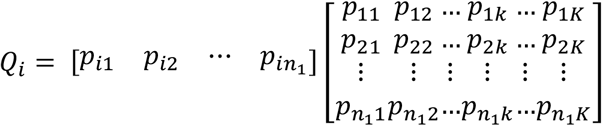

Finally, the contribution of each of the *N* input features in each of the *K* outputs can be written as

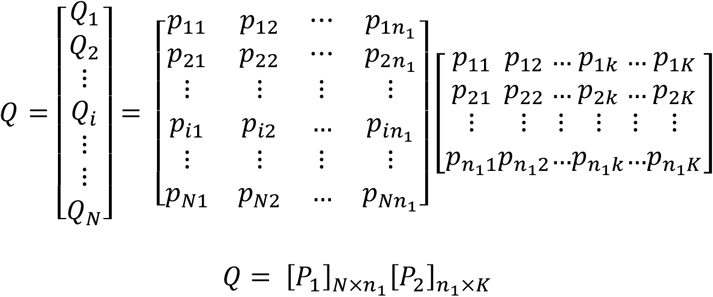

Case 2: Two hidden layers and K outputs

If we examine this problem closely, it becomes a recursion problem. If we start in the backward direction and take three layers together (two hidden and one output), it is the same as in Case 1, as discussed above.

Thus, the contribution of each of the *N*input features in each of the *K* outputs can be written as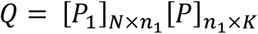

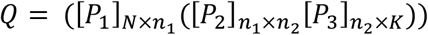

By using the commutative property of Matrices we can write *Q* as

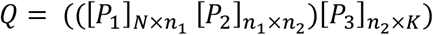

General case: (*L – 1*) hidden layers and K outputs

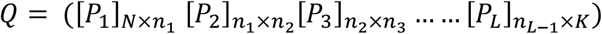

### Outlier feature detection

After estimating *Q*, we can make use of Mahalanobis distance to estimate the distance between each feature’s contribution vector from the average feature contribution vector. A larger distance for a feature indicates that the contributions made by that feature in the final prediction are higher than the average contribution made by the features.

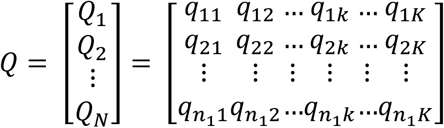

The Mahalanobis distance for *i*^*th*^ input feature can be defined as

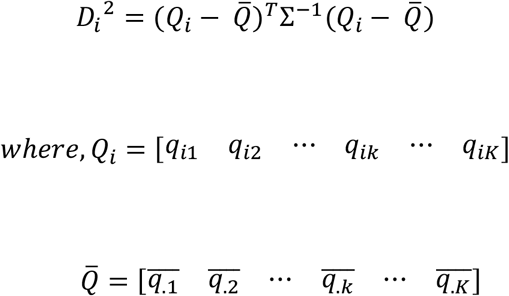

Σ is(*K* × *K*) covariance matrix of feature contributions

Mahalanobis distances are scaled by a constant λ known as genomic inflation factor, to obtain a test statistic that is approximated by a chi-square distribution with *K* degrees of freedom so that p-values can be estimated for each of the input features. Here, λ is defined as the ratio of the median of Mahalanobis distances to the median of the chi-square distribution with *K* degree of freedom (7).

The estimation of Mahalanobis distance relies on the main assumption that the estimated feature contributions follow a normal distribution. However, this assumption is often violated in Artificial Neural Networks (ANN) due to the utilization of regularization techniques like L1 and L2, leading to a positively skewed distribution of feature contributions. To address this issue, we propose a statistically sound approach. Our objective is to identify a theoretical distribution that best fits the estimated feature contributions. To achieve this, we fit a range of plausible distributions to the given data and estimate the parameters of each theoretical distribution. The selection of the most suitable statistical distribution is based on metrics such as likelihood, Akaike Information Criteria (AIC), and Bayesian Information Criteria (BIC). We then estimate the p-value using the fitted distribution and perform a Q-Q plot analysis of the estimated p-values. This helps us determine the threshold at which a deviation from the theoretical quantile occurs. Any features above this selected p-value threshold are considered plausible outlier features. By capturing the true nature of the feature contribution distribution, our proposed approach reduces the occurrence of false positives. For instance, when dealing with positively skewed data, plausible distributions to investigate could include the Beta distribution, Gamma distribution, Lognormal distribution, and Weibull distribution, among others.

## Discussion

We propose a robust and efficient framework for modelling the relationship between environmental factors and biological features using an ANN-based approach. We have focused on modelling all the output variables together instead of iterating over output variables and modelling each output sequentially, this significantly reduces the computational time. ANN are complex networks and are prone to overfitting if the defined architecture is not optimized. In this study, we focused on designing an architecture by performing hyperparameter optimization using random search, Hyperband, Bayesian optimization, and RandomSearch+lm. For each of the architectures obtained using the above method, we can monitor the loss of the training data and validation data over a fixed number of epochs and can choose the best architecture. Once the architecture is fixed, we can train the ANN on the entire dataset and measure the contribution of the input features in the prediction of output variables in the output layer. To estimate the contribution of each input feature in the output variables, we generalized a variable importance method that incorporates estimated weights from all layers in the ANN. The proposed feature selection method is efficient, as it uses a matrix multiplication approach to measure each input feature’s contribution in predicting each output variable in the output layer.

## Acknowledgements

We would like to express our sincere gratitude to Prof. Anil K. Ghosh from ISI, Kolkata, and Dr. Soham Sarkar from ISI, Delhi. Their expertise and insightful discussions during brainstorming sessions greatly enhanced the methodological development.

